# High-Throughput ChIPmentation: freely scalable, single day ChIPseq data generation from very low cell-numbers

**DOI:** 10.1101/426957

**Authors:** Charlotte Gustafsson, Ayla De Paepe, Christian Schmidl, Robert Månsson

## Abstract

Chromatin immunoprecipitation coupled to sequencing (ChIP-seq) is widely used to map histone modifications and transcription factor binding on a genome-wide level. Here, we present high-throughput ChIPmentation (HT-ChIPmentation) that eliminates the need for DNA purification prior to library amplification and reduces reverse-crosslinking time from hours to minutes. The resulting workflow is easily established, extremely rapid, and compatible with requirements for very low numbers of FACS sorted cells, high-throughput applications and single day data generation.

## BACKGROUND

The combination of chromatin immunoprecipitation with high-throughput sequencing (ChIP-seq) has become the method of choice for mapping chromatin-associated proteins and histone-modifications on a genome-wide level.

The ChIP-seq methodology has rapidly developed [1-4]. Despite this, performing ChIP-seq on limited cell-numbers and in a high-throughput manner remains technically challenging. This is largely due to decreasing input material leading to progressively increasing losses of material during DNA preparation and inefficiencies of enzymatic reactions used for library preparation. While elegant strategies have been developed to resolve these issues, they remain laborious and have not seen wider use [5-12].

ChIPmentation [3] effectively alleviates the issues associated with traditional library preparation methodologies by introducing sequencing-compatible adapters to bead-bound chromatin using Tn5 transposase (tagmentation). While fast and convenient, the methodology still relies on the use of traditional reverse crosslinking and DNA purification procedures prior to library amplification, hampering processing time, DNA recovery, and limiting scalability for high-throughput applications.

Here, we present freely scalable high-throughput ChIPmentation (HT-ChIPmentation) that by eliminating the need for DNA purification and traditional reverse-crosslinking prior to library amplification, dramatically reduces required time and input cell numbers. In comparison with current ChIP-seq variants [3, 5-12], HT-ChIPmentation is technically simple, extremely rapid and widely applicable, being compatible with both very low cell number requirements and high-throughput applications.

## RESULTS

The adapters introduced by Tn5 are covalently linked only to one strand of the tagmented DNA. The complete adapters, compatible with PCR amplification, are created through a subsequent extension reaction. With this in mind, we reasoned that performing adapter extension of tagmented bead-bound chromatin and high-temperature reverse crosslinking [6], would allow us to bypass the DNA purification step.

To validate this approach and benchmark it against standard ChIPmentation (Fig. 1A and Supplemental Fig. 1, Additional file 1), we FACS sorted defined numbers of formaldehyde fixed cells and performed ChIP with subsequent library preparation on cell numbers ranging from 0.1k-150k cells. HT-ChIPmentation indeed produced excellent sequencing profiles (Fig. 1B), and a consistent library size over >100-fold difference in input cell numbers (Supplemental Fig. 2A, Additional file 1).

**Figure 1.**
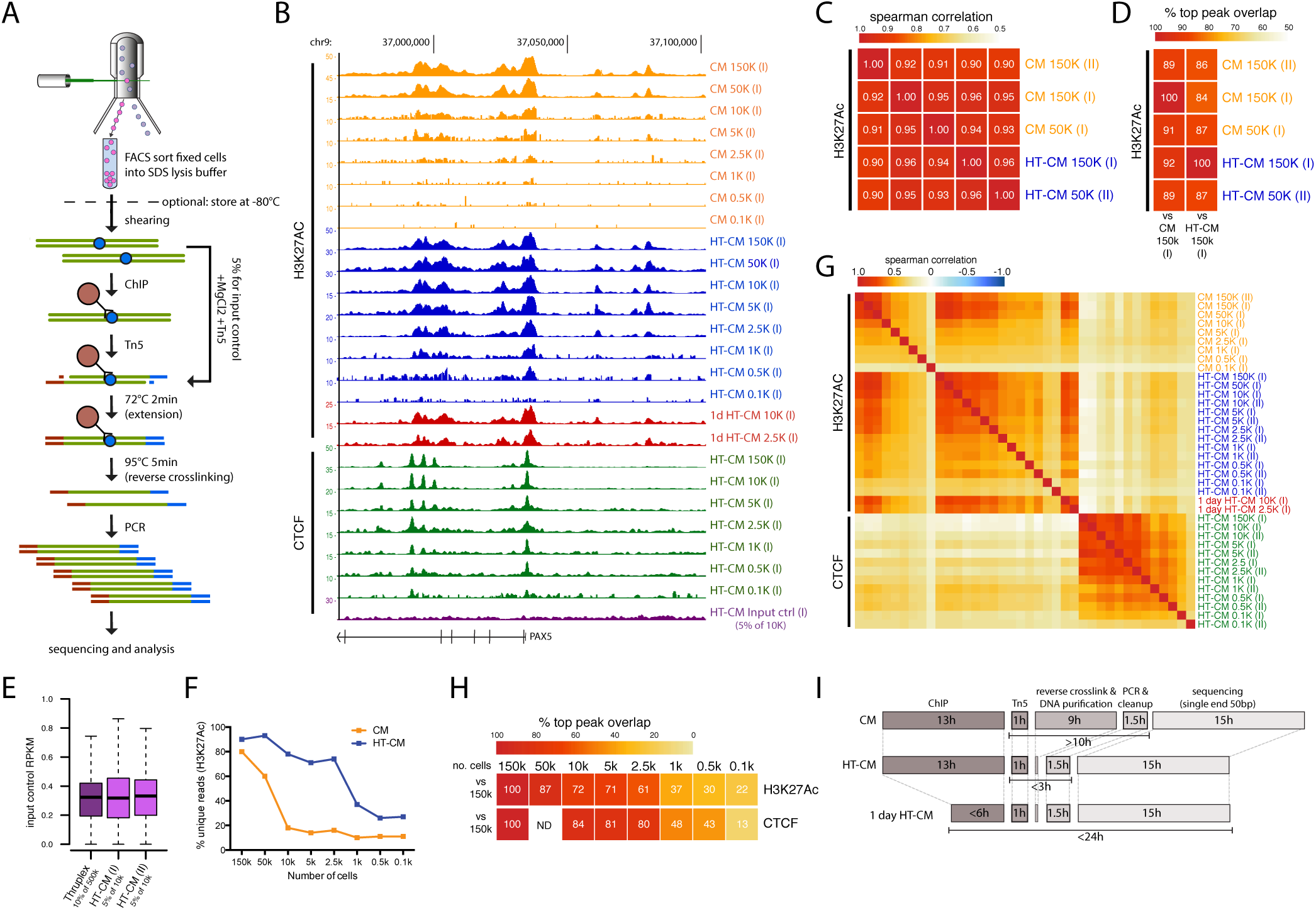
High-throughput ChIPmentation (HT-CMI through direct amplification of tagmented chromatin, allows for rapid and technically simple analysis of histone modifications and transcription factor binding in low numbers of FACS sorted cells. a) Schematic overview of the HT-CM workflow (for a direct comparison between the HT-CM and original CM methods, see Fig. S1, Additional file 1). In brief, FACS sorted cells are sonicated, subjected to ChIP and tagmented. Library amplification is subsequently done without prior DNA purification. Input controls are prepared through direct tagmentation of sonicated chromatin. b) Genome-browser profiles from CM, HT-CM and input control samples generated using indicated cell-numbers and antibodies. c) Correlation between H3K27Ac signals (in a merged catalog containing all peaks identified in displayed samples) generated using indicated methods and cell numbers. d) Overlap (%) between top peaks (peaks with the 50% highest peak quality scores) identified in high cell-number (150k and 50k) H3K27Ac HT-CM and CM samples. e) RPKM of 1kb bins covering the whole genome in input control samples generated using indicated method and cell-equivalents of chromatin. f) Percentage of unique reads in H3K27Ac HT-CM and CM samples generated in parallel. g) Correlation between H3K27Ac/CTCF signals in samples generated using indicated methods and cell-numbers. h) Overlap (%) between top peaks identified in H3K27Ac and CTCF HT-CM samples generated using indicated cell-numbers. ND, not done. i) Time required to perform ChIP, library preparation and sequencing for the CM, HT-CM and 1-day HT-CM workflows. Hours (h) needed to perform each step are indicated.

Looking specifically at H3K27Ac (a histone modification demarcating active promoters and enhancers [13]) HT-ChIPmentation and ChIPmentation samples generated in parallel from high cell-numbers (50k-150k cells), both methods generated high-quality data that is comparable in regard to: concordance of library profiles (Fig. 1B); mappability of sequencing reads (Supplemental Table 1, Additional file 1); correlation between samples (Fig. 1C); number, quality scores and signal range of identified peaks (Supplemental Fig. 2B-D, Additional file 1); and peak overlap (Fig. 1D).

To perform accurate peak calling, input controls were generated by direct tagmentation of 500 cell equivalents of sonicated chromatin (5% of 10k sonicated cells), subsequently processed in parallel with corresponding 10k HT-ChIPmentation samples (Fig. 1A). The HT-ChIPmentation compatible input controls produced similar results as input controls prepared using traditional library preparation methodology, in terms of library profiles and even genomic coverage (Fig. 1B and E).

We next compared H3K27Ac HT-ChIPmentation and ChIPmentation samples from progressively lower input cell-numbers. As expected, eliminating losses associated with DNA purification allowed HT-ChIPmentation samples to maintain much higher library complexity (>75% unique reads down to 2.5k cells) than ChIPmentation samples generated from the same number of cells (Fig. 1F). This difference in library quality was directly reflected in HT-ChIPmentation samples generated from a few thousand cells maintaining: consistent high quality library profiles (Fig. 1B); mappability (Supplemental Table 1, Additional file 1); number, quality scores and signal range of identified peaks (Supplemental Fig. 2B-D, Additional file 1); high correlation between samples (Fig. 1G); and high peak overlap (Fig. 1H). Similar results were obtained for H3K27Ac HT-ChIPmentation data generated in a single day (Fig. 1B, G and Supplemental figure 2B-D, Additional file 1). Based on the same metrics, CTCF (a chromatin organizing protein [14]) HT-ChIPmentation experiments further verified the robustness of the method with cell numbers in the range of a few thousands cells (Fig. 1B, G, H; Supplemental Fig. 2B-D and S3A-B, Additional file 1).

## DISCUSSION

Here we present HT-ChIPmentation, an improved and simplified tagmentation based approach to produce ChIP-seq libraries. We demonstrate that the adapters introduced by Tn5 can be extended directly on the bead-bound chromatin. Through this, we can combine ChIPmentation [3] with high-temperature reverse crosslinking and direct library amplification without prior DNA purification [6]. Even compared to the already technically simple and fast ChIPmentation method, HT-ChIPmentation is easier to perform and greatly reduces the time needed to produce sequencing ready libraries (Fig. 1I). In fact, HT-ChIPmentation together with sequencing can be performed in a single day (Fig. 1B, G and I; S2B-D, Additional file 1). This makes the protocol ideal for rapid data generation and compatible with the development of clinical diagnostic/prognostic applications relying on chromatin associated features to distinguish, for example, tumor subtypes [15, 16]

The removal of the DNA purification step, allows for fully taking advantage of that tagmentation of chromatin - as opposed to traditional adapter ligation [6, 8] - remains highly effective even with very limited input material ([3] and Supplemental Fig 2A, Additional file 1). Together, the reduced losses of material and effective addition of adapters, allows HT-ChIPmentation to be performed on just a few thousand FACS sorted cells with maintained quality and library complexity. Hence, HT-ChIPmentation provides a robust and technically simple workflow for characterizing epigenetic changes and transcription factor binding in rare subsets of cells.

Input controls are commonly used to exclude biases in the input material and as a negative control for identification of peak regions. Here we show that input controls can be prepared in parallel with HT-ChIPmentation samples, through direct tagmentation and library amplification of sonicated chromatin. The protocol requires very limited material (500 cell equivalents of sonicated chromatin), making it both feasible and convenient to directly prepare adequate controls for peak finding, also from rare subsets of cells.

The simplicity of the HT-ChIPmentation protocol - allowing for performing all steps from cells to amplified sequencing ready library without DNA purification - makes it perfectly suited for epigenetic characterization at any scale. While HT-ChIPmentation is directly compatible with full automation, experiments presented here were simply performed in 96-well plates using a multi-channel pipette, demonstrating that HT-ChIPmentation makes it highly feasible to perform epigenome scale projects in a matter of days using standard laboratory equipment.

## CONCLUSION

Here we introduce HT-ChIPmentation, an improved tagmentation based ChIP-seq protocol that through the extension of the Tn5`-i`nserted adapters on bead-bound chromatin, allows for direct library amplification without prior DNA purification. In comparison to current state-of-the-art ChIP-seq protocols [3, 5-12], HT-ChIPmentation is technically simple, extremely rapid and widely applicable, being compatible with very low cell number requirements, high-throughput applications and single day data generation. Taken together, HT-ChIPmentation provides a versatile and simplistic workflow attractive as the mainstay protocol for epigenome projects of any scale.

## METHODS

### Cells

Cultured MEC1 cells were stained with LIVE/DEAD fixable Aqua stain (Invitrogen) to allow for excluding cells dead already prior to fixation (during subsequent FACS sorting) and fixed using 1% PFA (Pierce). Aliquots of 10k cells were FACS sorted directly into 100μl SDS lysis buffer (50mM Tris/HCl, 0.5% SDS, and 10mM EDTA) supplemented with 1X cOmplete EDTA-free protease inhibitor (Roche) and stored at -80°C until use. For aliquots of cells (50k and 150k), where the sheath fluid volume is non-negligible, cells were sorted into PBS, spun down (2000g 5min) and resuspended in 100μl SDS lysis buffer prior to freezing. Sorting was performed using a BD FACSAriaIIu cell sorter (BD Biosciences) with an 85μm nozzle.

### Chromatin Immunoprecipitation and tagmentation

For ChIP, polyclonal anti-H3K27Ac (Diagenode, cat# C15410196, lot# A1723-0041D) antibody or anti-CTCF (Diagenode, cat# C15410210, lot# A2359-00234P) antibody was added to Protein G-coupled Dynabeads (ThermoFisher) in PBS with 0.5% BSA and incubated with rotation for 4h at 4°C (0.5h at RT for HT-ChIPmentation samples processed in a single day). For 50-150k cells, 10μl beads incubated with 3μg H3K27Ac or 1.5μg CTCF antibody were used per ChIP. For 0.1-10k cells, 2μl beads incubated with 0.6μg H3K27Ac or 0.3μg CTCF antibody were used per ChIP. Fixed cells (FACS sorted) frozen in SDS lysis buffer were thawed at room temperature. To perform ChIP on <10k cells, aliquots were diluted with SDS lysis buffer and 100μl containing the appropriate number of cells were processed. Cells were sonicated for 12 cycles of 30 sec on/30 sec off on high power using a Bioruptor Plus (Diagenode). To neutralize the SDS, Triton X100 was added to a final concentration of 1% along with 2μl 50x cOmplete protease inhibitor (final 1x). Samples were incubated at room temperature for 10min and when applicable 5% aliquots were saved for preparation of input controls. Antibody-coated Dynabeads were washed with PBS with 0.5% FCS and mixed with cell lysate in PCR tubes. Tubes were incubated rotating overnight (or 4h for HT-ChIPmentation samples processed in a single day) at 4°C.

Immunoprecipitated chromatin was washed with 150μ1 of low-salt buffer (50mM Tris/HCl, 150 mM NaCl, 0.1% SDS, 0.1% NaDOC, 1% Triton X-100, and 1mM EDTA), high-salt buffer (50mM Tris/HCl, 500mM NaCl, 0.1% SDS, 0.1% NaDoc, 1% Triton X-100, and 1 mM EDTA) and LiCl buffer (10 mM Tris/HCl, 250 mM LiCl, 0.5% IGEPAL CA-630, 0.5% NaDOC, and 1mM EDTA), followed by two washes with TE buffer (10mM Tris/HCl and 1mM EDTA) and two washes with ice cold Tris/HCl pH8. For tagmentation, bead bound chromatin was resuspended in 30μl of tagmentation buffer, 1μl of transposase (Nextera, Illumina) was added and samples were incubated at 37°C for 10 minutes followed by two washes with low-salt buffer.

### High-throughput ChIPmentation library preparation

For HT-CM samples, bead bound tagmented chromatin was diluted in 20μl of water. PCR master mix (Nextera, Illumina) and indexed amplification primers [17] (0.125uM final concentration) was added and libraries prepared using the following program: 72°C 5min (adapter extension); 95°C 5min (reverse cross-linking); followed by 11 cycles of 98°C 10s, 63°C 30s and 72°C 3min.

For preparation of HT-CM compatible input controls, 1μl of 50mM MgCh was added to 5μl sonicated lysate (5% aliquot of 10k samples) to neutralize the EDTA in the SDS lysis buffer. 30μl of tagmentation buffer and 1μl transposase (Nextera, Illumina) was added, and samples were incubated at 37°C for 10min. 22.5μl of the transposition reaction were combined with 15μl of PCR master mix and 2.5μl of primer mix (Nextera, Illumina). Libraries were subsequently amplified as described for HT-ChIPmentation samples.

### ChIPmentation library preparation

For standard reverse crosslinking, chromatin complexes were diluted with 200μl ChIP elution buffer (10mM Tris/HCl, 0.5% SDS, 300mM NaCl, and 5 mM EDTA) and 2μl of 20μg/ml proteinase K (Thermo Scientific). Samples were vortexed and incubated with shaking overnight at 65°C. After reverse crosslinking, 1μl 20μg/ml RNase (Sigma) was added and incubated at 37°C for 30min. After another 2h of incubation with 2μl of proteinase K (20mg/ml) at 55°C, samples were placed in a magnet to trap magnetic beads and supernatants were collected. DNA purification was carried out using Qiagen MinElute PCR Purification Kit. 15μl of PCR master mix and 5μl of primer mix (Nextera, Illumina) was added to 20μl of eluted DNA, and libraries were amplified as described for HT-ChIPmentation libraries.

### Preparation of conventional input control

Sonicated material from 50k cells was reverse crosslinked as described for ChIPmentation. 2ng of DNA was used for library preparation using the ThruPLEX DNA-seq kit (Rubicon Genomics) with 11 cycles of PCR amplification.

### Post-PCR library cleanup and sequencing

After PCR amplification, library cleanup was done using Agencourt AmPureXP beads (Beckman Coulter) at a ratio of 1:1. DNA concentrations in purified samples were measured using the Qubit dsDNA HS Kit (Invitrogen). Libraries were pooled and single-end sequenced (50 cycles) using the Nextseq500 platform (Illumina).

### Basic processing of ChIP-seq and input control sequencing data

Quality of the sequenced samples was assessed using `FastQC` v0.11.5 [18]. Samples were mapped to the human reference genome (hg19) using Bowtie2 v2.2.3 [19] with default settings. Further basic processing was performed using HOMER v4.8.3 [20]. Specifically, mapped reads were converted into tagdirectories by the `makeTagDirectory` command using settings for the human genome (`-genome hg19`) and removing duplicate reads by allowing only one tag to start per base pair (`-tbp 1`).

### Genome Browser visualizations

Bedgraphs were created for each sample using HOMER's `makeUCSCfile`. Tracks were uploaded and visualized using the UCSC genome browser [21].

### Peak finding and plotting peak metrics

Peak finding was performed using the `findPeaks` command in HOMER. Peaks were called using default settings for histone modifications (`-style histone`) and transcription factors (`-style factor`) for H3K27Ac and CTCF respectively with input (`-i`) as a control. Visualization was done in R v3.1.0 [22], using the built in `barplot` and `barplot` R-functions to plot peak numbers and peak quality scores, respectively.

### Making and annotating peak catalogs

Peak catalogs were created by merging all peak files of samples analyzed using HOMER's `mergePeaks` command. Setting used (`-size given`) ensured that peaks with literal overlap were merged to one peak while peaks unique to one sample were directly added to the peak catalog. Subsequently, peak catalogs were annotated with unnormalized (`-raw`) read counts within peaks in the catalog for each individual sample using HOMER's `annotatePeaks.pl` script.

### Plotting peak read distributions and correlation between samples

Raw counts were log normalized in R as follows: `log(df[,countsCols]+1,2)`. Log2 counts were subsequently plotted using the build in `barplot`R-function. These same Log2 counts were used to calculate sample correlations, using the build-in `cor` R-function with spearman correlation. Correlation matrices were visualized with the `pheatmap` function from the pheatmap R-package using color scales generated with the build-in `colorRampPalette` R-function.

### Plotting reads within 1kb bins for input control samples

A file containing 1kb bins covering the whole genome was created using the `makewindows` command from bedtools v2.26.0 [23] using a window size of 1kb (`-w 1000`). Chromosome sizes were retrieved as follows: `mysql — user=genome --host=genome-mysql.cse.ucsc.edu-A -e"select chrom, size from hg19.chromInfo" > hg19.genome`. Raw reads in each 1kb bin for each input control were counted using HOMER's `annotatePeaks.p1` script, as described above. Raw read distributions were converted to RPKM in R based on the standard RPKM formula. Resulting RPKM distributions were plotted with the build-in `barplot` R-function.

### Determining top peak overlap

Peaks identified in individual samples were overlapped with in-house code using the IRanges [24] R-package. Top peaks overlap was considered to be the percentage of high quality peaks (50% of peaks with highest quality scores) in the reference sample that overlap (≥1 bp) with a peak in the second sample. For purposes of determining peak overlap, CTCF peaks were extended with 50bp up and downstream, considering `findPeaks` with `-style factor` only calls a small region around the peak maximum. Peak overlaps were visualized using the `pheatmap` function from the pheatmap R-package using color scales generated with the build-in `colorRampPalette` R-function.

### Comparing library complexity

To compare duplication rates between HT-ChIPmentation and ChIPmentation samples, fastq files were randomly down-sampled to the total number of reads in the smallest file for each cell number. Down sampling was performed using the `fastq-sample` script from fastq-tools v0.8 [25]. Fraction of unique reads was subsequently determined for each file using `FastQC` v0.11.5.

### Motif enrichment analysis

Enrichments of known transcription factor binding motifs in peaks were identified using HOMER’s `findMotifsGenome.pl` script with default settings.

## DECLARATIONS

### Ethics approval and consent to participate

Not applicable.

### Consent for publication

Not applicable.

### Availability of data and materials

The generated data sets are available from the European Nucleotide Archive [26] under the study accession number: PRJEB23059

### Competing interests

C.S. has a pending patent application for ChIPmentation. C.G., A.D.P., and R.M. have no competing financial interests.

## Funding

This work was funded by the Swedish Cancer Foundation (Cancerfonden), the Swedish Research Council (VR), the Knut and Alice Wallenberg Foundation (KAW) and the Swedish Foundation for Strategic Research (SSF).

## Authors’ contributions

C.G. and R.M. devised the HT-ChIPmentation workflow and planned the study; C.G. performed experiments; C.G., A.D.P. and R.M. analyzed the data; C.S. provided critical insights; R.M. supervised the research; and all authors contributed to the writing of the manuscript.

## Acknowledgements

We thank: Prof. Joakim Dillner and colleagues for access to the NextSeq 500 system; Prof. Anders Rosén for providing the MEC1 cell-line; Dr. Yin C. Lin for critical reading of the manuscript; and UPPMAX Next Generation Sequencing Cluster & Storage (UPPNEX) for computational resources.

